## Supplemental Figs. 1-6 for "Oligodendrocyte-derived IL-33 functions as a microglial survival factor during neuroinvasive flavivirus infection"

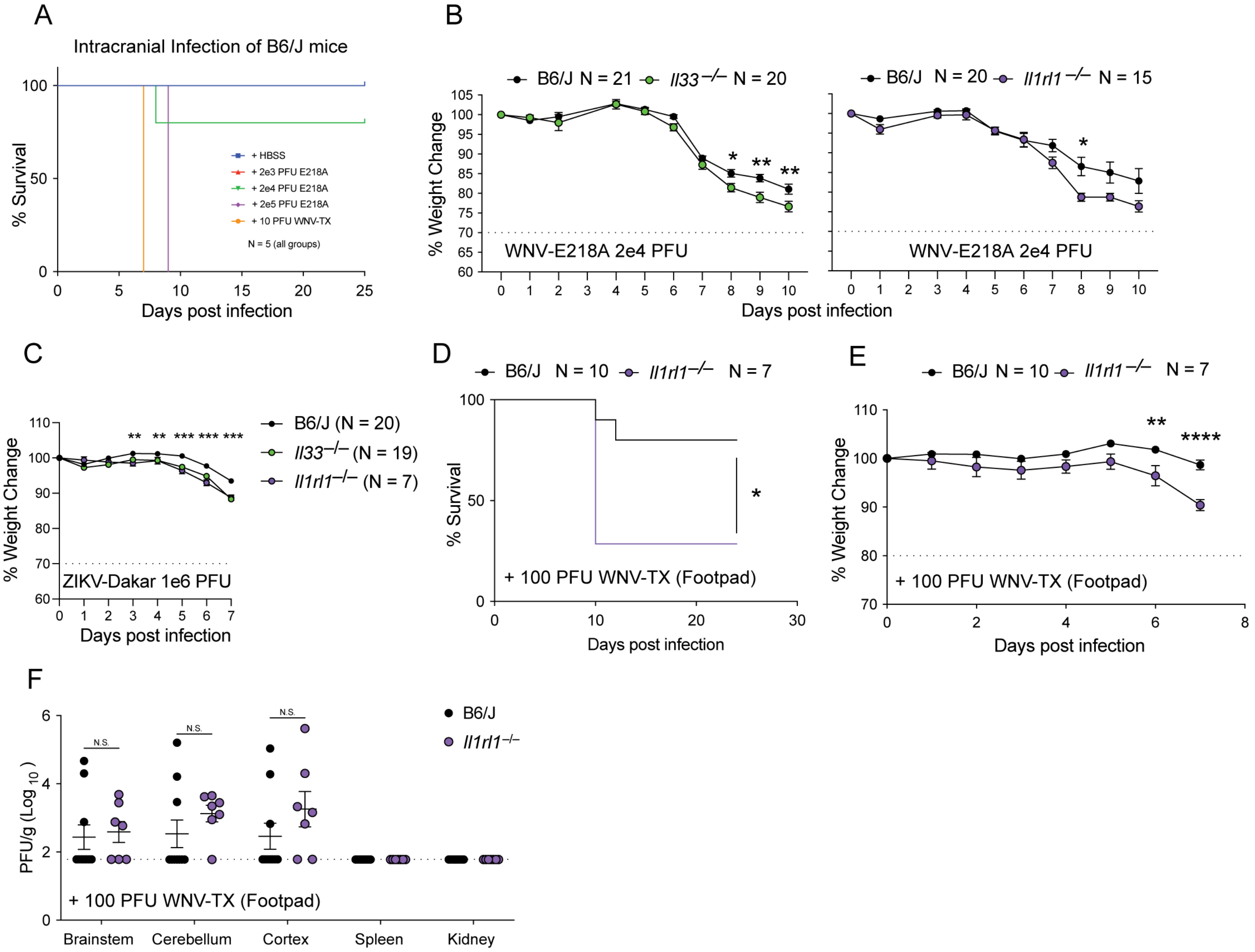


Figure S1 (related to Figure 1). The IL-33 pathway improves disease outcomes across multiple models of flavivirus infection

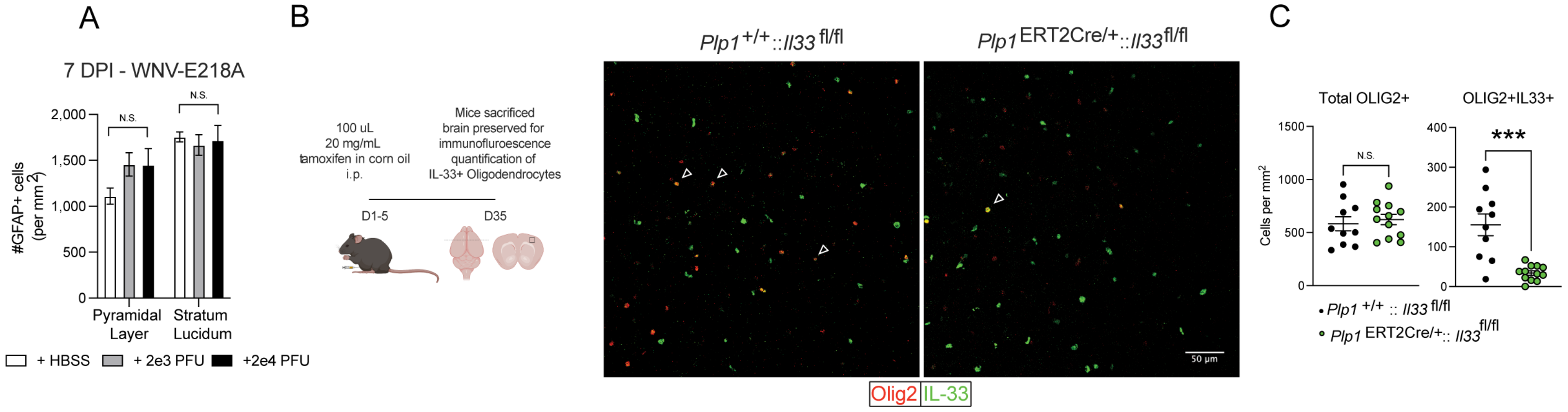


Figure S2 (related to Figure 2). Deletion of IL-33 in oligodendrocytes with tamoxifen treatment of *Plp1*$^{ERT2Cre/+}$::*Il33*$^{fl/fl}$ mice

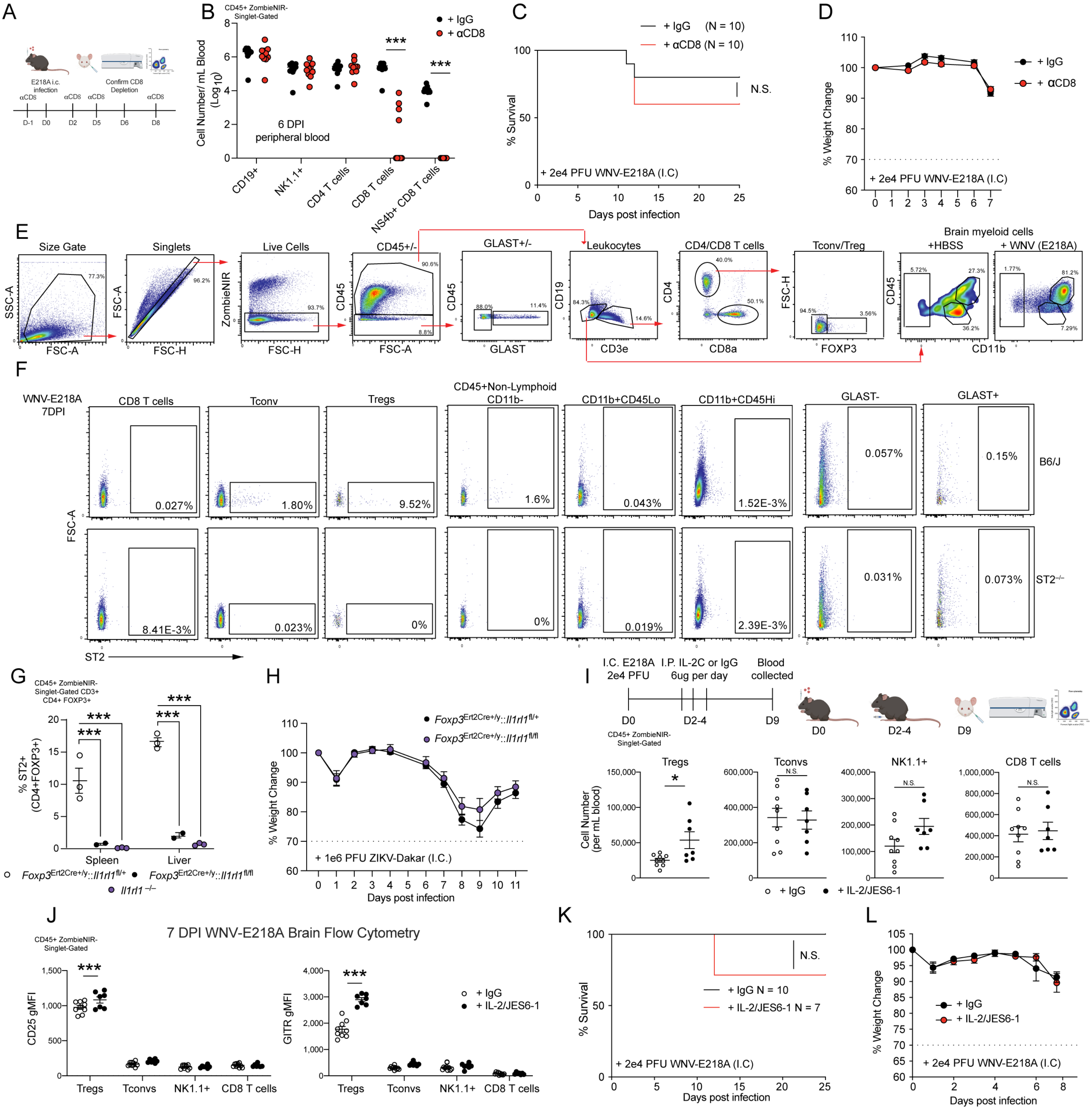


Figure S3 (related to Figure 3). ST2 is highly expressed on adaptive immune subsets, but this expression is dispensible for host survival.

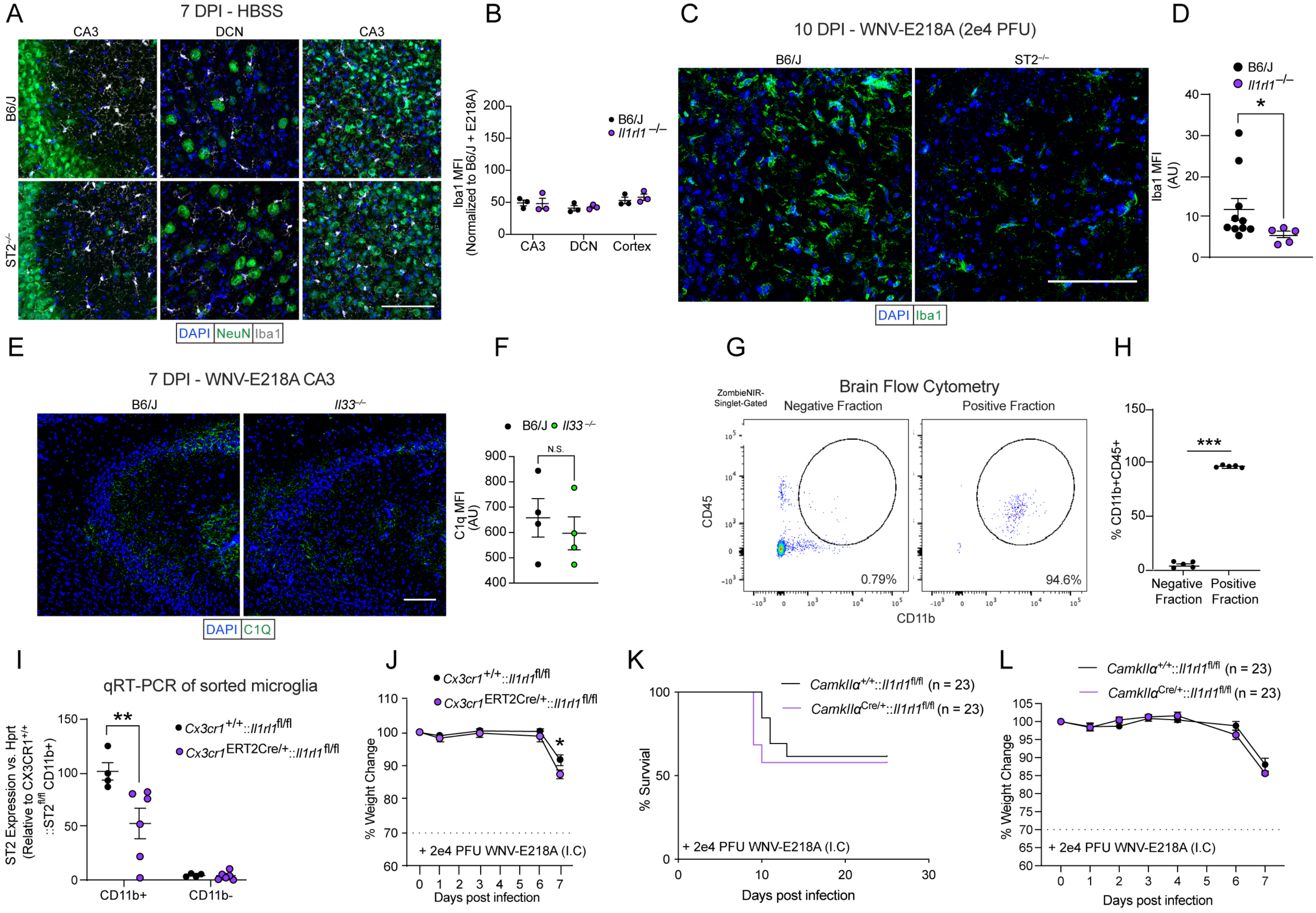


Figure S4 (related to Figure 4). Phenotypes of ST2 signaling on microglial activation during CNS Flavivirus infection.

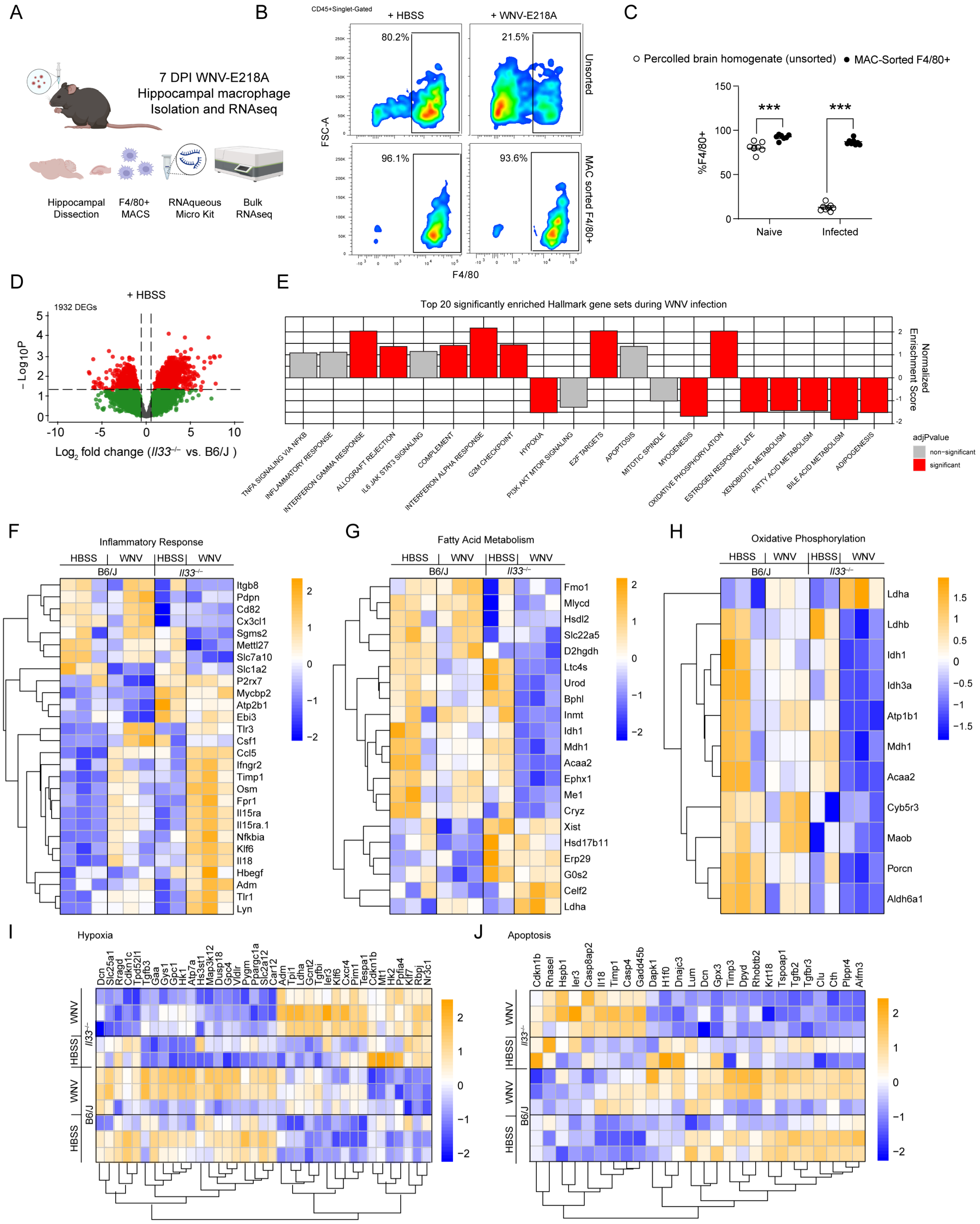


Figure S5 (related to Figure 5). Hallmark pathways enriched in *Il33*−/− hippocampal macrophages during WNV infection.

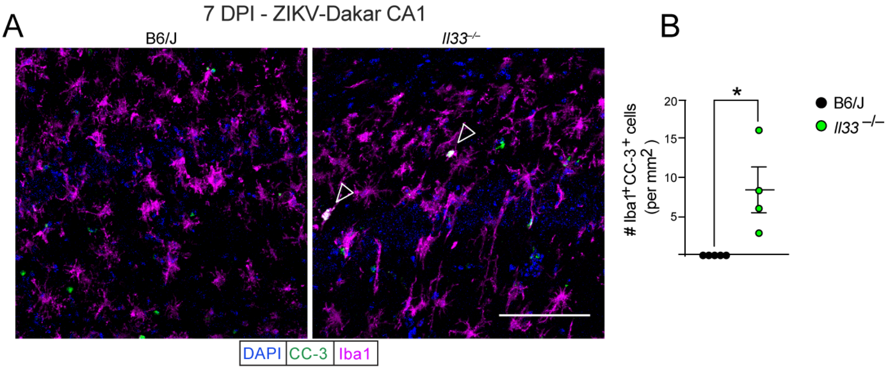


Figure S6 (related to Figure 6). IL-33 signaling prevents brain macrophage apoptosis following intracranial ZIKV-Dakar infection.
